# Disruption of ultrasonic vocalizations with systemic administration of the non-competitive N-methyl-D-aspartate receptor antagonist MK-801 in adult male mice

**DOI:** 10.1101/2024.08.25.609588

**Authors:** Ryuto Tamura, Mizuki Yamamoto, Yusuke Ujihara, Haruki Kasahara, Shuntaro Matsushima, Daiki Nasukawa, Kazuko Hayashi, Kota Yamada, Masashi Tanaka, Koji Toda

## Abstract

**Objective:** Social behavior and communication, including ultrasonic vocalizations (USVs), are important for survival and reproductive success across species. N-methyl-D-aspartate (NMDA) receptors contribute to the neural mechanisms underlying social behavior, and pharmacological modulation of these receptors can alter both social behavior and USVs in rodents. However, the effects of NMDA receptor blockade on dyadic USVs and social approach during male-female interactions in mice remain incompletely understood. We therefore investigated the effects of systemic administration of MK-801, a noncompetitive NMDA receptor antagonist, on dyadic vocal activity, social approach, locomotor behavior, and excretory outcomes in adult male mice.

**Methods:** Sexually experienced male mice were tested in male-female interaction and same-sex social-preference tasks. Using VocalMat, a machine-learning-based tool, we classified the USVs recorded during male-female interactions into 11 call categories and examined their temporal relationship to the proximity of the male subject to the female stimulus.

**Results:** Administration of MK-801 to the male dose-dependently reduced the total number of USVs recorded from the male-female dyad, with a marked reduction at the highest dose, while leaving the treated male’s proximity to the female stimulus unchanged. In the same-sex social-preference task, MK-801 reduced the time spent near the social stimulus at the highest dose, although preference for the social stimulus over the non-social object remained significant under all treatment conditions. MK-801 also increased locomotor activity and reduced fecal output and estimated urine output in the open-field task. These findings demonstrate that systemic NMDA receptor blockade differentially affects dyadic vocal communication, social approach, locomotor activity, and excretory outcomes.

**Conclusion:** Systemic MK-801 administration to the male reduced the number of USVs recorded during male–female interactions and produced distinct effects on social approach, locomotor activity, and excretory outcomes. These findings indicate that NMDA receptor blockade differentially affects multiple behavioral domains, although the mechanisms underlying the reduction in USVs remain to be determined.

**Significant Outcomes:**

- Systemic MK-801 administration to male mice dose-dependently reduced the number of ultrasonic vocalizations recorded from male-female dyads without significantly altering the treated male’s proximity to the female stimulus.
- In the same-sex social-preference task, the highest dose of MK-801 reduced the time spent near the social stimulus, while preference for the social stimulus over the non-social object was preserved under all treatment conditions.
- MK-801 increased locomotor activity and center occupancy in the open-field task while reducing fecal output and estimated urine output, demonstrating dissociable effects across vocal, social, locomotor, and excretory outcomes.

**Limitations:**

- The single-microphone recording system did not permit individual ultrasonic vocalizations to be assigned to either member of the male–female dyad.
- Because MK-801 was administered systemically, the study could not identify the neural, peripheral, respiratory, or vocal-motor mechanisms underlying the reduction in USVs.
- The study used only C57BL/6J mice, and the estrous stage of the stimulus females was not determined, limiting assessment of strain-, species-, sex-, and reproductive-status-related variability.

## Introduction

N-methyl-D-aspartate (NMDA) receptors play fundamental roles in activity-dependent synaptic plasticity and cognitive processes.^1^ Reduced NMDA receptor signaling has been implicated in several neuropsychiatric conditions, including schizophrenia,^2–4^ in which alterations in social and communicative functioning have also been described.^5,6^ These clinical and neurobiological observations have contributed to broader interest in the behavioral consequences of NMDA receptor hypofunction. Pharmacological blockade of NMDA receptors provides a controlled approach for examining these consequences in experimental animals.

MK-801 (dizocilpine) is a noncompetitive NMDA receptor antagonist widely used to induce experimentally controlled NMDA receptor hypofunction in rodents. Systemic MK-801 administration impairs learning and memory,^7–16^ disrupts sensorimotor gating,^17^ increases locomotor activity and stereotyped behavior,^18–23^ and alters social interaction and social approach.^21,24–27^ The magnitude and direction of these social effects can vary with dose, species, and testing conditions. A social-preference task therefore provides a useful measure for characterizing how NMDA receptor blockade affects approach toward a conspecific. However, whether NMDA receptor blockade also alters vocal activity during social encounters, and whether such changes occur together with alterations in social approach, remain incompletely understood. Because systemic administration cannot identify the neural or peripheral site of drug action, changes across behavioral measures should not be assumed to arise from a single underlying mechanism.

Vocal behavior provides a quantifiable component of social interaction that complements conventional measures of social approach. In adult mice, USVs are observed predominantly during specific social encounters, most prominently in male– female interactions, although they have also been reported in male–male and female– female contexts.^28,29^ In contrast to the well-characterized drug-induced 50-kHz USVs of rats, amphetamine- and amphetamine-associated cue-induced USVs were not detected in repeatedly treated C57BL/6J mice.^30^ During male–female encounters, USVs are commonly recorded in association with investigation, courtship, and reproductive behavior. Male USVs have been described as a representative precopulatory measure of sexual motivation, alongside other measures such as urine marking, approach toward a receptive female, and preference for females or female-derived odors.^31^ Studies testing males individually have shown that exposure to an intact female or freshly collected female urine can elicit USVs, although urine-evoked vocalizations are generally less robust than those recorded during direct male–female interactions and may depend on stimulus conditions, including urine storage.^28,32–34^ Experimental comparisons of vocalizing and devocalized males and playback experiments have shown that females approach or spend more time near vocalizing males or males paired with USV playback.^35–37^ Courtship vocal activity has also been reported to correlate with reproductive outcomes in laboratory and wild-derived house mice, including the number of deliveries and the latency to the first litter, although the limited available evidence does not establish a causal relationship.^36–38^ Moreover, males are generally the primary vocalizers during male–female encounters, although sound-source localization has demonstrated that females can also contribute a minority of USVs during freely moving courtship interactions.^39^ Consequently, the vocalizing animal cannot be determined with certainty from recordings obtained without caller-identification methods.

Recent methodological advances have enabled increasingly accurate detection, segmentation, classification, sound-source localization, and quantification of rodent USVs.^39–43^ These approaches make it possible to examine vocal activity alongside the spatial and temporal organization of social behavior. However, dyadic vocal recordings and social-approach measures provide related but distinct information. A change in the total number of USVs recorded from a dyad may occur without a corresponding change in proximity, whereas altered approach behavior may influence vocal activity indirectly. Simultaneous assessment of these measures is therefore important for distinguishing changes in dyadic vocal activity from broader alterations in social engagement. Measurement of locomotor activity and other behavioral outcomes is also necessary because MK-801-induced hyperactivity or stereotypy may complicate the interpretation of social and vocal measures.

Evidence from rats indicates that glutamatergic NMDA receptor signaling contributes to the modulation of USVs across several experimental contexts. Rats selectively bred for low levels of play-induced 50-kHz USVs show molecular alterations involving the NMDA receptor family, providing evidence of an association between NMDA receptor-related mechanisms and vocal phenotype.^44^ More directly, infusion of MK-801 into the nucleus accumbens core reduced the increase in 50-kHz USVs induced by repeated cocaine administration in rats.^45^ Systemic MK-801 has also been shown to modify the acute, sensitized, and conditioned effects of amphetamine on rat 50-kHz USVs.^46^ These findings support a contribution of NMDA receptor signaling to the regulation of rat USVs but also indicate that its effects depend on the neural site, pharmacological manipulation, and behavioral context. Whether NMDA receptor blockade similarly alters USVs recorded during male-female interactions in mice, and how such changes relate to social-approach behavior, remain incompletely understood. Moreover, because the effects of systemic MK-801 may involve social, vocal-motor, respiratory, autonomic, and locomotor processes, dyadic USV recordings alone cannot identify the mechanism underlying any observed change.

The present study examined the effects of acute systemic MK-801 administration to adult male mice across complementary behavioral domains. First, we classified USVs recorded during male-female interactions and examined their temporal relationship to the proximity of the treated male to the female stimulus. Second, we determined whether MK-801 administration to the male altered the number of USVs recorded from the dyad and the male’s approach behavior toward the female. Third, we used a same-sex social-preference task to examine whether MK-801 affected approach toward a novel conspecific in a context not centered on courtship or vocal output, thereby determining whether its effects extended to a broader measure of sociability. Finally, we evaluated locomotor activity and excretory outcomes in an open-field task to characterize additional behavioral and physiological effects relevant to the interpretation of the social and vocal findings. By integrating these complementary measures, we aimed to characterize the effects of systemic NMDA receptor blockade on dyadic vocal activity and social behavior while recognizing that systemic pharmacological administration does not permit the underlying neural or peripheral mechanisms to be localized.

## Material and methods

### Subjects

Adult male wild-type C57BL/6J mice were used as subject animals. Subject males were 3–6 months old in Experiments 1 and 2 and 2–4 months old in Experiment 3. The female stimulus mice used in Experiment 1 were 2–3 months old. All animals were maintained on a 12-h light/12-h dark cycle, and all experiments were conducted during the dark phase. Food and water were available ad libitum, and the body weights of the subject mice were monitored daily. All animal procedures were approved by the Animal Research Committee of Keio University. The study is reported in accordance with the ARRIVE guidelines (https://arriveguidelines.org).

### Drug

The NMDA receptor antagonist (+)-MK-801 hydrogen maleate (MK-801; Sigma-Aldrich, St Louis, Missouri, USA) was dissolved in saline, divided into small aliquots, and stored in a refrigerator at 4°C until further use. Prior to all the experiments, we administered the MK-801 solution intraperitoneally at a dose of 0.1, 0.2, or 0.4 mg/kg. For the habituation and control conditions, saline was administered intraperitoneally, as in the drug condition. The doses of MK-801 used in this study (0.1, 0.2, and 0.4 mg/kg) were selected based on previous studies in mice and rats.^10–13^

### Experimental procedure

#### Experiment 1: Social interaction task

Experiment 1 included seven adult male C57BL/6J mice and was designed to examine the effects of MK-801 administration to the male on USVs recorded during male-female interactions and on the male’s approach behavior toward a female stimulus (Figure 1A). The social-interaction apparatus and general recording configuration were based on those described previously.^47,48^ Before behavioral testing, each subject male was housed with a female mouse for 1 week to provide prior heterosexual experience. The males were subsequently habituated to the experimental apparatus, a custom-made transparent vinyl chloride box (28 cm long × 8.5 cm wide × 30 cm high), for at least 3 days. Testing was conducted on 4 days per week for 3 consecutive weeks. Because preliminary observations indicated day-to-day variation in the number of recorded USVs, particularly on the first testing day of each week, practice sessions were conducted on the first and fourth days. In each week, a saline control session was conducted on the second day, followed by an MK-801 session on the third day. Thus, each male underwent three saline sessions, each performed on the day immediately preceding the corresponding MK-801 session, and one session at each MK-801 dose (0.1, 0.2, and 0.4 mg/kg). One MK-801 dose was tested per week, and the order of the three doses across the three testing weeks was randomized across animals. For each outcome measure, the three saline-session values were averaged within each animal to obtain a single subject-level saline value for statistical analysis and graphical presentation. Thus, each animal contributed one value to each of the four treatment conditions: mean saline and MK-801 at 0.1, 0.2, and 0.4 mg/kg. Saline or MK-801 was administered intraperitoneally 5 min before the start of the corresponding behavioral session. Each session consisted of habituation and social-interaction phases. During the 15-min habituation phase, the subject male was allowed to explore the apparatus freely in the presence of an empty cylindrical basket. The male was then temporarily removed, and a novel female stimulus mouse was placed inside the basket (8 cm in diameter × 10 cm high) located at one end of the apparatus. A different female stimulus was used for each test session. Because the female remained confined within the basket, direct physical contact, mating, and pregnancy were prevented. A 500-mL empty glass bottle was placed on top of the basket to prevent the animals from climbing onto it. The subject male was then returned to the apparatus and allowed to explore freely for 10 min. A single ultrasonic microphone (CM16/CMPA, Avisoft Bioacoustics, Germany) was positioned above the apparatus, and USVs were recorded throughout the social-interaction phase. Audio signals were digitized through an audio interface (UR824, Steinberg, Japan) at a sampling rate of 192 kHz with 16-bit resolution and stored on a computer for subsequent analysis. A video camera (C980GR, Logicool, Japan) positioned above the apparatus recorded the location and movement of the subject male. Testing was conducted inside a sound-attenuating chamber (ENV-018V, Med Associates Inc., USA) to minimize external noise. Because a single overhead microphone was used and the recording system did not permit sound-source localization, individual calls could not be assigned to either animal. USVs were therefore quantified at the dyad level and are referred to throughout the manuscript as USVs recorded during male–female interactions or from the male–female dyad. After each session, the inside surfaces of the apparatus, basket, and bottle were cleaned with 70% ethanol and allowed to air-dry for 30 min.

**Figure 1.**
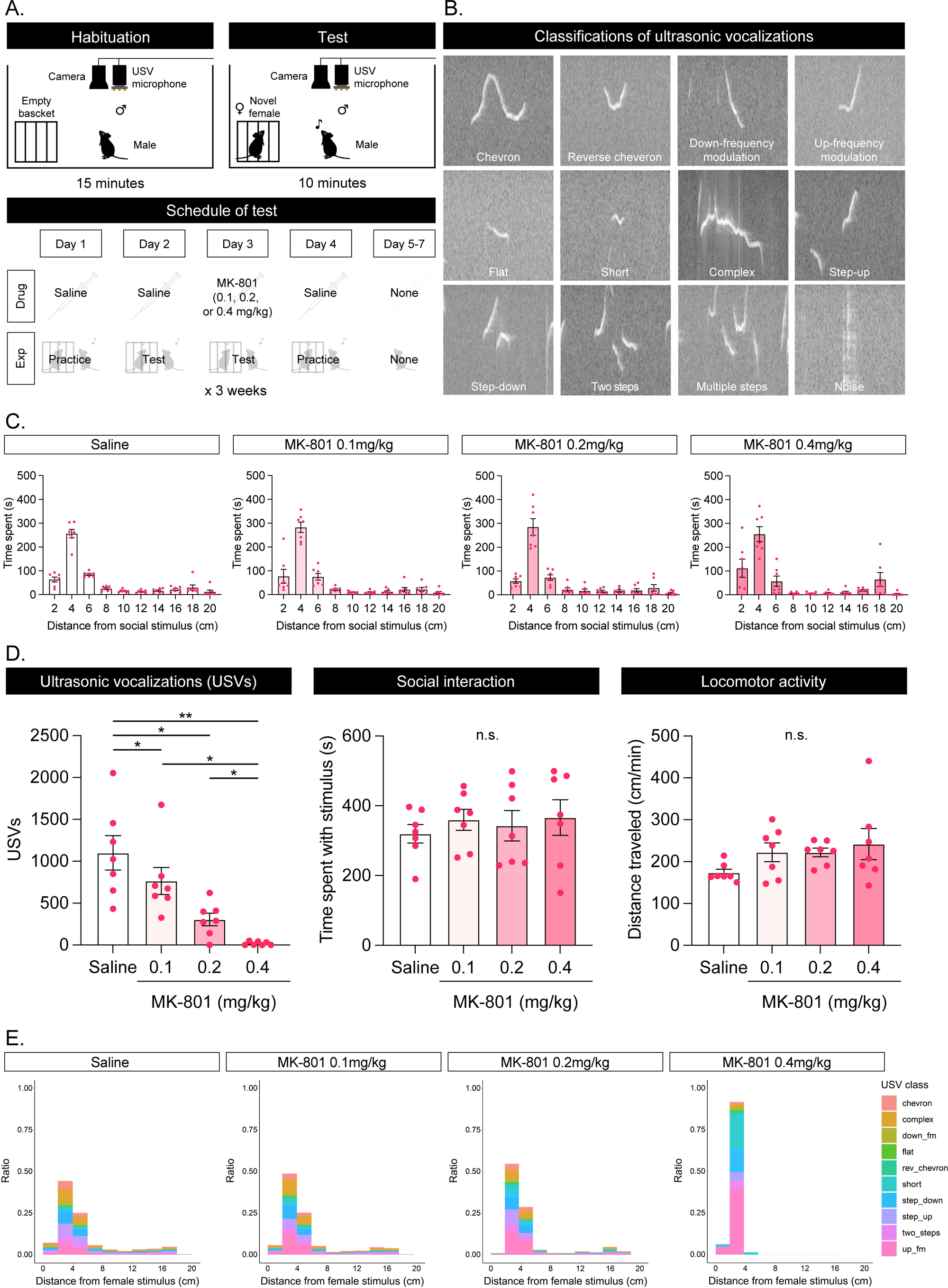
Effects of MK-801 administration to the male on dyadic USVs and behavior in the social-interaction task. **A.** The apparatus used for the test was a custom-made transparent vinyl chloride box (28 cm long × 8.5 cm wide × 30 cm high), with a basket containing a novel female stimulus mouse placed at one end of the box. Subject males were intraperitoneally injected with saline or MK-801 (0.1, 0.2, or 0.4 mg/kg) 5 min before being placed in the apparatus. After a 15-min habituation period without the female stimulus (upper left panel), the subject male was temporarily removed from the apparatus, and a novel female stimulus mouse was placed in the basket. The subject male was then returned to the apparatus and allowed to explore freely for 10 min (upper right panel). Experiments were conducted in a sound-attenuating chamber to minimize external noise. Testing was conducted on 4 days per week for 3 consecutive weeks (bottom panel). Given the day-to-day variation in the number of recorded USVs, practice sessions were conducted on the first and fourth days of each week. Saline and MK-801 sessions were conducted on the second and third days, respectively, so that each MK-801 session was preceded by a saline session. Each male therefore underwent three saline sessions and one session at each MK-801 dose. For statistical analysis and graphical presentation, the values from the three saline sessions were averaged within each animal to generate a single subject-level saline value. **B.** USVs were classified into 11 categories—chevron, reverse chevron, downward frequency modulation, upward frequency modulation, flat, short, complex, step-up, step-down, two-step, and multiple steps—and distinguished from noise using VocalMat. **C.** Time spent by the treated male at different distances from the female stimulus mouse. **D.** Total number of USVs recorded from the dyad (left panel), time spent by the treated male near the female stimulus (center panel), and locomotor activity of the treated male (right panel). **E.** Distribution of classified USVs according to the distance of the treated male from the female stimulus. In panels C and D, each data point represents one animal. For the saline condition, each data point represents the mean of the three saline-session values obtained from that animal. Error bars indicate the standard error of the mean. \**p* < 0.05 and \*\**p* < 0.01. *n* = 7 animals.

#### Experiment 2: Social preference task

To examine the effects of MK-801 on social preference, we used a procedure adapted from those described previously.^48,49^ Thirty-two adult male C57BL/6J mice (3–6 months old) were used as subjects, and two adult male C57BL/6J mice (3 months old) were used as social stimuli (Figure 2A). All mice were group-housed with two to four animals per home cage. A mixed design was used, with treatment condition as a between-subjects factor and stimulus type as a within-subject factor; each subject mouse was assigned to only one treatment condition and was tested once. The apparatus was a custom-made white vinyl chloride box measuring 50 cm long × 25 cm wide × 30 cm high. Two white cylindrical baskets (8 cm in diameter × 10 cm high) were positioned at opposite ends of the apparatus. During the test phase, a novel male C57BL/6J stimulus mouse was confined within one basket, and a novel object was placed in the other basket. A 500-mL empty glass bottle was placed on top of each basket to prevent the subject mouse from climbing onto it. Two partition walls (5 cm long × 5 mm thick × 30 cm high) were positioned on either side of the center of the apparatus to partially separate the two stimulus areas. A video camera (C920r, Logicool, Japan) was positioned 80 cm above the floor of the apparatus, and behavior was recorded on a Windows computer. Subject mice were randomly assigned to receive an intraperitoneal injection of saline or MK-801 at 0.1, 0.2, or 0.4 mg/kg. Five minutes after injection, each subject was placed in the apparatus and allowed to habituate for 15 min in the presence of two empty baskets. The subject was then temporarily removed, after which a novel male stimulus mouse and a novel object were placed in separate baskets at opposite ends of the apparatus. The subject was returned to the center of the apparatus and allowed to explore freely for 10 min. White noise (75 dB) was presented throughout the experiment to minimize interference from external sounds. After each session, the apparatus, baskets, and bottles were cleaned with 70% ethanol and allowed to air-dry for 30 min.

**Figure 2.**
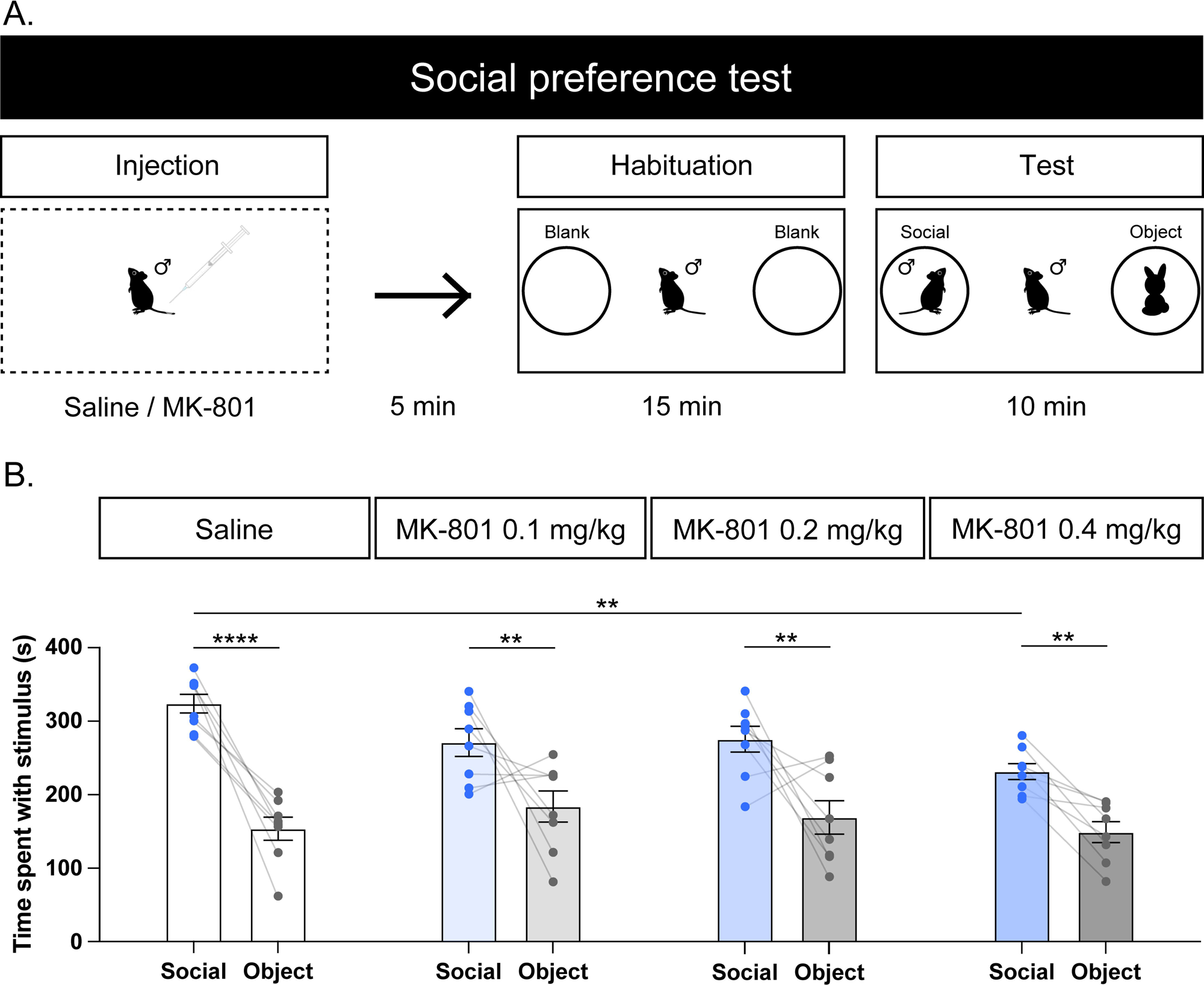
Effects of MK-801 on social preference toward a novel same-sex conspecific. **A.** Schematic representation of the social-preference task. The apparatus was a custom-made white vinyl chloride box (50 cm long × 25 cm wide × 30 cm high) with two cylindrical baskets positioned at opposite ends. Subject mice were intraperitoneally injected with saline or MK-801 (0.1, 0.2, or 0.4 mg/kg) and placed in the apparatus 5 min after injection. After a 15-min habituation period with two empty baskets, the subject mouse was temporarily removed. A novel male C57BL/6J stimulus mouse and a novel object were then placed in separate baskets at opposite ends of the apparatus. The subject mouse was returned to the center of the apparatus and allowed to explore freely for 10 min. **B.** Time spent near the novel male social stimulus and the novel object under each treatment condition. A subject mouse was defined as being near a stimulus when its center-of-mass position was located within 7.5 cm of the edge of the basket containing that stimulus. Error bars indicate the standard error of the mean. \*\**p* < 0.01 and \*\*\*\**p* < 0.0001. *n* = 8 animals per group.

#### Experiment 3: Open-field task

To examine the effects of MK-801 on spontaneous locomotor activity, we conducted an open-field task using 11 adult male C57BL/6J mice (2–4 months old). The apparatus and experimental procedure were adapted from those described in previous studies.^48,50–52^ Habituation to the apparatus and saline administration were performed for 60 min/day for 3 days before the experiment. The apparatus consisted of a custom-made white vinyl chloride box measuring 50 cm long × 50 cm wide × 50 cm high. A camera (Logicool HD Webcam C920r, Logicool) was positioned 106 cm above the floor of the box, and videos were recorded on a Windows PC. During habituation, the animals were placed in the open-field apparatus immediately after receiving an intraperitoneal injection of saline and were allowed to explore freely for 60 min. The order of saline and MK-801 administration (0.1, 0.2, and 0.4 mg/kg) was randomized, with an interval of more than 2 days between sessions to minimize carryover effects of MK-801. Each test session lasted 60 min. White noise (75 dB) was presented throughout the experiment to mask external noise. Immediately after completion of each open-field session, the entire floor surface of the apparatus was systematically wiped with pre-weighed filter paper to absorb the urine deposited during the test. The filter paper was weighed again after urine absorption, and the difference between the pre- and post-collection weights was calculated as an estimate of urine output and expressed in grams. Because this procedure may not have recovered all urine and may have been affected by evaporation, the measurement was regarded as a semi-quantitative estimate rather than an exact determination of urine volume. After urine and fecal samples had been collected, the box was wiped with 70% alcohol and allowed to air-dry for 30 min.

### Analysis

For all experiments, data were analyzed using RStudio (version 2022.02.0; RStudio PBC, MA, USA) and GraphPad Prism (version 10.3.0; GraphPad Software, CA, USA). In Experiment 1, VocalMat, an open-source USV analysis framework, was used to detect and classify USVs.^41^ The starting times of the video and audio recordings were synchronized using a custom-built circuit comprising a button, an LED, and a small speaker. In all experiments, the position and movement of the subject mouse were tracked from video recordings using Bonsai, an open-source visual programming framework for computer vision analysis.^53^ Video frames were converted to grayscale, smoothed, and converted into inverted binary images, and the mouse was identified by applying a contrast threshold. Locomotor activity was quantified from changes in the center-of-mass coordinates across successive video frames. In Experiment 1, the position of each subject male was tracked using its center-of-mass coordinate. For the analysis of social proximity, the male was defined as being near the female stimulus when its center of mass was located within 4.0 cm of the edge of the stimulus basket. Separately, to visualize the spatial distributions of occupancy and USV occurrence in Figure 1C and E, the distance between the male’s center of mass and the stimulus basket was divided into 2-cm bins. For each recorded USV, the corresponding distance bin was determined from the position of the treated male at the time the call was detected; this analysis does not identify the location or identity of the vocalizing animal. The relatively small amount of data in the innermost 0–2-cm bin reflects a limitation of center-of-mass–based tracking rather than an absence of close investigation. Because the center of mass is located near the torso, it rarely enters this innermost bin unless the mouse stands upright or leans toward the basket. Thus, even when the mouse approaches the basket with its nose, its center of mass typically remains outside the 0– 2-cm bin. In Experiment 2, the subject mouse was defined as being near a stimulus when its center of mass was located within 7.5 cm of the edge of the basket containing that stimulus. In Experiment 3, the center area was defined as the central 25 cm × 25 cm portion of the open-field apparatus.

For the statistical analysis of Experiment 1, the values obtained during the three saline sessions were averaged within each animal to generate a single subject-level saline value for each outcome. Thus, each animal contributed one value to each of the four treatment conditions: mean saline and MK-801 at 0.1, 0.2, and 0.4 mg/kg. The effects of treatment on the total number of USVs recorded from the dyad, locomotor activity of the treated male, and time spent by the treated male near the female stimulus were analyzed using one-way repeated-measures analysis of variance (ANOVA), with treatment condition as the within-subject factor. When a significant main effect of treatment was detected, pairwise comparisons were conducted using Tukey’s multiple-comparisons test. The Greenhouse–Geisser correction was applied when the assumption of sphericity was not met, and adjusted degrees of freedom are reported where applicable. In Experiment 2, the time spent near each stimulus was analyzed using a two-way mixed-design ANOVA, with treatment condition (saline and MK-801 at 0.1, 0.2, and 0.4 mg/kg) as the between-subjects factor and stimulus type (social stimulus and non-social object) as the within-subject factor. Sidak’s multiple-comparisons test was used for post hoc comparisons. Statistical tests for Experiment 3 were selected according to the repeated-measures design and are reported with the corresponding results. All tests were two-tailed, statistical significance was defined as *p* < 0.05, and data are presented as the mean ± SEM.

## Results

USVs recorded during male–female interactions were classified into 11 categories using VocalMat (Figure 1B). Across treatment conditions, USVs were recorded predominantly during periods when the treated male was near the female stimulus (Figure 1C). MK-801 dose-dependently reduced the total number of USVs recorded from the male– female dyad (Figure 1D, left panel; one-way repeated-measures ANOVA with the Greenhouse-Geisser correction, *F*(1.424, 8.546) = 19.86, *p* = 0.0010). Tukey’s multiple-comparisons test showed that the number of USVs was significantly lower following MK-801 administration at 0.1 mg/kg (adjusted *p* < 0.05), 0.2 mg/kg (adjusted *p* < 0.05), and 0.4 mg/kg (adjusted *p* < 0.01) than in the mean saline condition. The number of USVs was also significantly lower following administration of 0.4 mg/kg MK-801 than following administration of either 0.1 mg/kg (adjusted *p* < 0.05) or 0.2 mg/kg MK-801 (adjusted *p* < 0.05). Very few calls were detected at the highest dose of MK-801. MK-801 did not significantly affect either the time that the treated male spent near the female stimulus (Figure 1D, center panel; one-way repeated-measures ANOVA with the Greenhouse-Geisser correction, *F*(1.759, 10.55) = 0.370, *p* = 0.6735) or the locomotor activity of the treated male (Figure 1D, right panel; one-way repeated-measures ANOVA with the Greenhouse-Geisser correction, *F*(1.192, 7.150) = 1.328, *p* = 0.2970). Descriptive examination of the data presented in 2-cm distance bins did not reveal a consistent relationship between individual USV categories and the distance of the treated male from the female stimulus (Figure 1E). Because the recording system did not permit sound-source localization, the vocalizing animal could not be identified.

To examine the effects of intraperitoneal MK-801 administration on preference for a novel same-sex social stimulus, we used a social-preference task (Figure 2). Mice received an intraperitoneal injection of saline or MK-801 at 0.1, 0.2, or 0.4 mg/kg and were placed in the apparatus 5 min after injection. Following a 15-min habituation period with two empty baskets, the mice were temporarily removed from the apparatus. A novel male C57BL/6J stimulus mouse and a novel object were then placed in separate baskets at opposite ends of the apparatus. The subject mouse was returned to the center of the apparatus and allowed to explore freely for 10 min (Figure 2A).

The time spent near each stimulus was analyzed using a two-way mixed-design ANOVA, with treatment condition as the between-subjects factor and stimulus type as the within-subject factor. The analysis revealed significant main effects of stimulus type (*F*(1, 28) = 60.98, *p* < 0.0001) and treatment (*F*(3, 28) = 5.636, *p* = 0.0038), but no significant treatment × stimulus-type interaction (*F*(3, 28) = 1.701, *p* = 0.1894; Figure 2B). Sidak’s multiple-comparisons test showed that mice spent significantly more time near the social stimulus than near the non-social object under all treatment conditions (all adjusted *p* < 0.01), indicating that social preference was preserved following MK-801 administration. However, mice administered 0.4 mg/kg MK-801 spent significantly less time near the social stimulus than saline-treated mice (adjusted *p* = 0.0021). Thus, the highest dose of MK-801 reduced social approach without significantly altering the relative preference for the social stimulus over the non-social object.

To examine the effects of MK-801 on locomotor activity and excretory outcomes, we conducted an open-field task (Figure 3A). Following three daily 60-min habituation sessions, mice were intraperitoneally injected with saline or MK-801 (0.1, 0.2, or 0.4 mg/kg) and placed in the apparatus 5 min after injection. MK-801 significantly affected locomotor activity (Figure 3B, left panel; *F*(2.177, 21.77) = 7.817, *p* = 0.0023, repeated-measures one-way ANOVA). Locomotor activity was significantly higher following all three doses of MK-801 than following saline (saline vs. 0.1 mg/kg MK-801, *p* < 0.0001; saline vs. 0.2 mg/kg MK-801, *p* = 0.0184; saline vs. 0.4 mg/kg MK-801, *p* = 0.0367; Tukey’s post hoc tests). Time-course analysis showed that locomotor activity gradually decreased over time under the saline condition. In contrast, this decline was absent following MK-801 administration, and locomotor activity tended to increase, particularly in the 0.2 and 0.4 mg/kg conditions. The effect became apparent approximately 15 min after injection and remained evident until the end of the 60-min observation period (Figure 3B, right panel).

**Figure 3.**
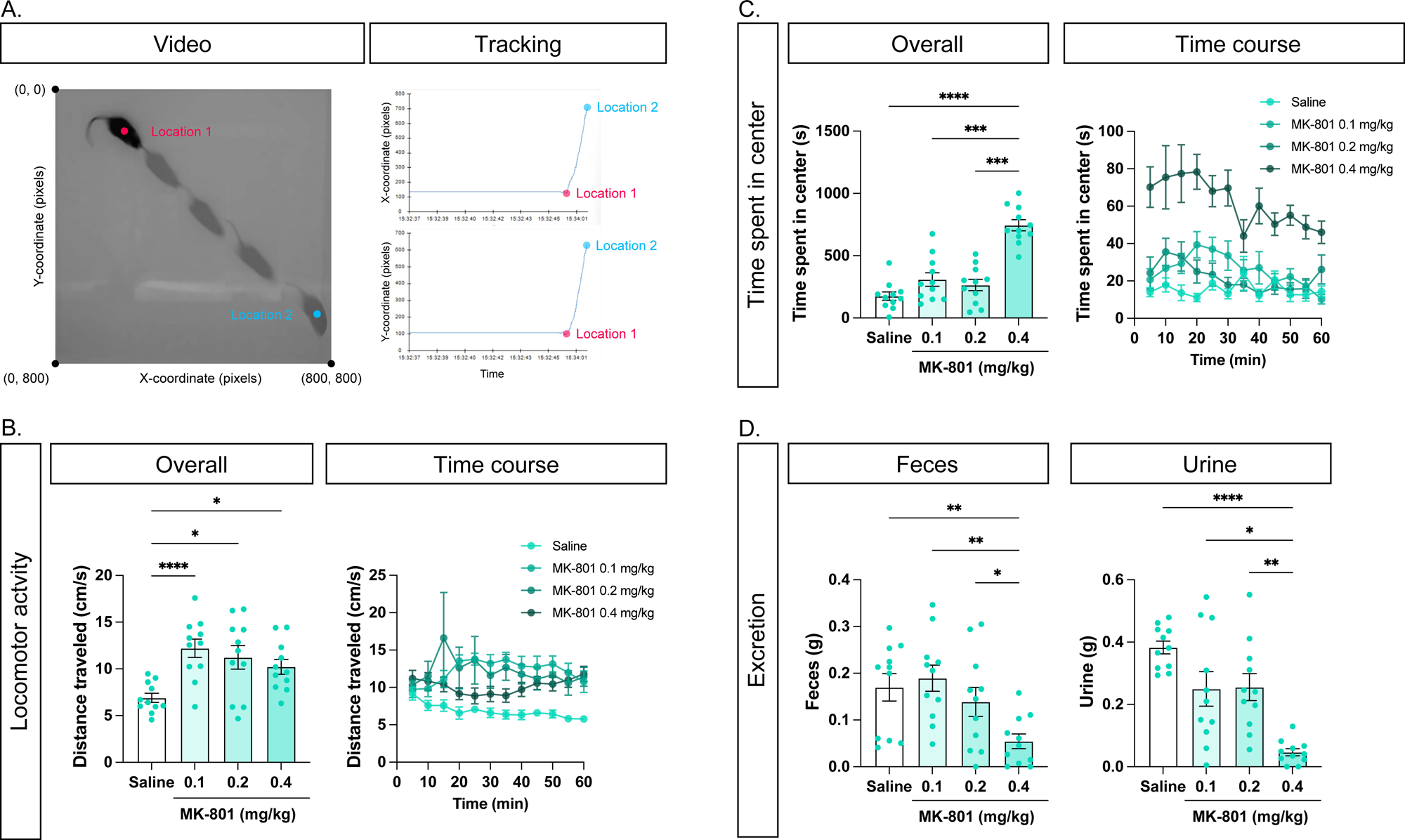
Effects of MK-801 on the open-field task. **A.** Example of the tracking procedure. **B.** Overall locomotor activity from 10 to 60 min after the start of the open-field task following administration of saline or MK-801 (left panel). Time course of locomotor activity (right panel). **C.** Time spent in the center of the open-field apparatus from 10 to 60 min after the start of the task (left panel). Time course of center occupancy (right panel). The center area was defined as the central 25 cm × 25 cm portion of the apparatus. **D.** Fecal output measured after the 60-min open-field task (left panel) and estimated urine output measured after the task (right panel). Error bars indicate the standard error of the mean. \**p* < 0.05, \*\**p* < 0.01, \*\*\**p* < 0.001, and \*\*\*\**p* < 0.0001. *n* = 11 animals.

We next examined the time that the mice spent in the center of the open-field apparatus. MK-801 significantly affected center occupancy (Figure 3C, left panel; *F*(2.573, 25.73) = 37.42, *p* < 0.0001, repeated-measures one-way ANOVA). Mice spent significantly more time in the center following administration of 0.4 mg/kg MK-801 than following saline (*p* < 0.0001), 0.1 mg/kg MK-801 (*p* = 0.0003), or 0.2 mg/kg MK-801 (*p* = 0.0002; Tukey’s post hoc tests). Time-course analysis showed that the increase in center occupancy emerged shortly after MK-801 administration and persisted throughout the 60-min observation period (Figure 3C, right panel).

To assess excretory outcomes that may be influenced by autonomic, enteric, behavioral, and peripheral mechanisms, fecal output and estimated urine output were quantified after each open-field session. MK-801 significantly affected both fecal output (Figure 3D, left panel; *F*(2.086, 20.86) = 5.555, *p* = 0.0109, repeated-measures one-way ANOVA) and estimated urine output (Figure 3D, right panel; *F*(1.740, 17.40) = 12.82, *p* = 0.0006, repeated-measures one-way ANOVA). Both measures decreased with increasing doses of MK-801. Fecal output was significantly lower following administration of 0.4 mg/kg MK-801 than following saline (*p* = 0.0093), 0.1 mg/kg MK-801 (*p* = 0.0090), or 0.2 mg/kg MK-801 (*p* = 0.0224). Estimated urine output was also significantly lower following administration of 0.4 mg/kg MK-801 than following saline (*p* < 0.0001), 0.1 mg/kg MK-801 (*p* = 0.0196), or 0.2 mg/kg MK-801 (*p* = 0.0024; Tukey’s post hoc tests).

## Discussion

The present study examined the effects of acute systemic MK-801 administration to adult male mice across dyadic vocal, social, locomotor, and excretory measures. Administration of MK-801 to the male reduced the number of USVs recorded from male-female dyads but did not significantly alter the time that the treated male spent near the female stimulus. In the same-sex social-preference task, the highest dose of MK-801 reduced the time spent near the social stimulus, although the mice continued to spend significantly more time near the social stimulus than near the non-social object under all treatment conditions. In the open-field task, MK-801 increased locomotor activity and center occupancy while reducing fecal output and estimated urine output. These findings demonstrate that systemic NMDA receptor blockade produces dissociable changes across dyadic vocal activity, social approach, locomotor behavior, and excretory outcomes.

We classified the USVs recorded during male–female interactions into 11 categories and distinguished them from non-vocal sounds using VocalMat.^41^ The calls were recorded predominantly during periods when the treated male was near the female stimulus, indicating that vocal activity was temporally associated with the male– female interaction. Previous studies have established that males are the primary vocalizers during male–female encounters in mice.^28,54^ Sound-source localization in freely interacting C57BL/6J mice confirmed that both males and females can vocalize during courtship but attributed the large majority of localized vocalizations to males.^39^ Hammerschmidt et al.^55^ demonstrated that male and female C57BL/6J mice produce acoustically similar USVs in resident–intruder paradigms and that the resident animal was the primary vocalizer when paired with an anesthetized intruder; however, that study did not directly examine vocal exchanges between males and females. Matsumoto et al.^56^ similarly identified the male as the vocalizing animal in the male– female context in C57BL/6J mice, whereas context-dependent reciprocal vocalizations were observed in ICR mice. The available evidence indicates that the USVs recorded in the present study were most likely produced predominantly by the male. Administration of MK-801 to the male reduced the total number of USVs recorded during the interaction, with very few calls detected at the highest dose (0.4 mg/kg). The most straightforward interpretation is therefore that MK-801 suppressed vocal production by the treated male. Nevertheless, because our single-microphone recording system did not permit caller identification under the specific experimental condition in which the male explored a female confined within a basket, a contribution from the female cannot be completely excluded. We also found no consistent relationship between individual USV categories and the distance of the treated male from the female stimulus. Although several classification systems have been proposed for mouse USVs,^57–59^ the biological significance of individual call categories remains incompletely understood and is likely to depend on social and behavioral context.^60^ Future studies combining methods that permit caller identification, such as sound-source localization or individually mounted microphones,^39,56,61^ with markerless pose estimation^62–64^ and temporally resolved classification of behavioral dynamics^65,66^ will be important for confirming the identity of the vocalizing animal and clarifying how individual call categories relate to ongoing social behavior.

Evidence from rats indicates that glutamatergic NMDA receptor signaling contributes to the modulation of USVs across several experimental contexts. Complementary genetic evidence has been obtained from rats selectively bred for low levels of play-induced 50-kHz USVs, in which transcriptomic network analysis identified the NMDA receptor family as a molecular hub associated with the low-vocalization phenotype.^44^ Pharmacological studies have provided more direct evidence. Sohn et al.^45^ showed that infusion of MK-801 into the nucleus accumbens core reduced the increase in 50-kHz USVs induced by repeated cocaine administration, implicating local NMDA receptor signaling in drug-induced vocal activity. Costa et al.^46^ similarly demonstrated that systemic MK-801 modified the acute, sensitized, and conditioned effects of amphetamine on rat 50-kHz USVs and was associated with altered Zif-268 expression in the nucleus accumbens, dorsolateral striatum, and medial prefrontal cortex. These findings support a role for NMDA receptor signaling in the modulation of rat USVs but also indicate that the effects depend on the neural site, pharmacological manipulation, and behavioral context. Previous rat studies have further shown that drug-induced changes in USVs can diverge from those in locomotor activity, conditioned place preference, and other behavioral outcomes.^46,67–69^ Nevertheless, differences in species, social context, call type, and pharmacological protocol limit direct extrapolation from rat 50-kHz USVs to the dyadic USVs recorded from mice in the present study.

The mechanisms underlying the reduction in dyadic USVs require cautious interpretation. In Experiment 1, MK-801 administration to the male reduced the number of recorded USVs without significantly altering the time that the treated male spent near the female stimulus. Thus, the reduction in vocal activity should not be interpreted simply as evidence of a primary deficit in social approach. Because mouse USV production depends on the coordination of expiration, laryngeal motor activity, and supralaryngeal vocal mechanisms, MK-801 may have altered respiratory or vocal-motor output rather than, or in addition to, social communication processes.^70,71^ The absence of a reduction in locomotor activity in Experiment 1 argues against generalized motor suppression as the sole explanation for the decrease in dyadic USVs. Moreover, the increase in locomotor activity observed in the open-field task provides additional evidence against a generalized suppression of motor behavior. Nevertheless, these findings do not exclude disruption of the specific respiratory and motor patterns required for vocalization. The combination of increased locomotor activity and center occupancy with reduced excretory outcomes in the open-field task also cannot be explained by a single autonomic mechanism. In particular, increased center occupancy should not be interpreted as an anxiolytic-like effect without accounting for MK-801-induced hyperactivity and stereotyped movement. Because respiration, laryngeal activity, autonomic function, and stereotyped behavior were not measured directly, the present experiments cannot distinguish among changes in social communication, respiratory or vocal-motor control, autonomic regulation, and indirect changes in the behavior of the untreated female. The results are therefore most conservatively interpreted as demonstrating dissociable effects of systemic NMDA receptor blockade across dyadic vocal activity, social approach, locomotor behavior, and excretory outcomes rather than identifying a single mechanism that accounts for all observed changes.

Although cortical regions may modulate vocal behavior, the cerebral cortex does not appear to be indispensable for the basic production of mouse USVs. Mice lacking the hippocampus and large portions of the cerebral cortex developed broadly preserved pup isolation calls and adult courtship USVs, with only minor differences in adult call-type usage.^72^ This finding indicates that subcortical and brainstem circuits can support the fundamental generation of mouse USVs and argues against attributing the reduction observed after systemic MK-801 administration specifically to disruption of cortical vocal circuits. Nevertheless, cortical regions may modulate the initiation, timing, or acoustic organization of vocalizations according to behavioral and social context. In rats, the association between MK-801-induced changes in 50-kHz USVs and altered Zif-268 expression in the medial prefrontal cortex reported by Costa et al.^46^ suggests possible involvement of prefrontal circuitry but does not establish the medial prefrontal cortex as the direct site of drug action. In mice, the anterior cingulate cortex has been implicated in the modulation of social vocalizations.^73^ In marmosets, neonatal lesions of the anterior cingulate cortex altered the subsequent development and social use of vocalizations.^74^ Other studies in humans and nonhuman primates have implicated anterior cingulate and ventrolateral prefrontal regions in vocal control.^75–78^ At the subcortical level, lesion evidence from cats has historically implicated the midbrain periaqueductal gray in vocal production,^79^ while circuit-level studies in mice have identified the periaqueductal gray as a key component of the descending circuitry controlling social vocalizations.^80,81^ Because MK-801 was administered systemically in the present study, our findings cannot distinguish among cortical modulation, subcortical or brainstem vocal mechanisms, respiratory or vocal-motor effects, and indirect behavioral effects within the dyad. Region- and cell type-specific manipulation of NMDA receptor signaling, combined with caller-identification and physiological measurements, will be required to determine the mechanisms underlying the reduction in dyadic USVs.

The effects of MK-801 on social approach differed between the two social-behavior experiments. In Experiment 1, MK-801 administration to the male did not significantly alter the time spent near the female stimulus, whereas the highest dose of MK-801 reduced the time spent near the same-sex social stimulus in Experiment 2. Nevertheless, the treatment × stimulus-type interaction was not significant in Experiment 2, and the mice continued to spend significantly more time near the social stimulus than near the non-social object under all treatment conditions. Thus, the highest dose reduced overall social approach without eliminating the relative preference for the social stimulus. The difference between the experiments may reflect the sex of the stimulus animal, the motivational context, or the experimental apparatus. In particular, Experiment 1 used a narrow chamber designed to facilitate simultaneous audio and video recording, which may have restricted the range of possible social-approach behaviors and reduced the sensitivity of proximity duration as a measure of sociability. Previous studies examining pharmacological or genetic disruption of NMDA receptor function have also reported variable effects on rodent social behavior.^21,24,25,82,83^ Such variability may arise from differences in dose, treatment schedule, apparatus, habituation, social context, and the potentially confounding effects of NMDA receptor blockade on locomotion, learning, memory, and social recognition. Future studies using well-controlled but more naturalistic social-interaction paradigms will be necessary to clarify how NMDA receptor hypofunction affects distinct components of social behavior.

In the open-field task, MK-801 increased locomotor activity and the time spent in the center of the apparatus. Previous studies have shown that MK-801 can induce stereotyped behaviors, including circling and jumping, in a dose-dependent manner.^9,23^ Such stereotyped movements may have contributed both to the increase in center occupancy and to the less pronounced increase in total locomotor activity at 0.4 mg/kg than at lower doses. However, because stereotyped movements were not quantified separately in the present study, their contribution remains uncertain. Future studies incorporating detailed classification of movement patterns will be required to distinguish hyperlocomotion and stereotypy from anxiety-related changes in spatial exploration.

MK-801 also dose-dependently reduced fecal output and estimated urine output in the open-field task. The effects of acute MK-801 administration on these excretory measures have not previously been extensively characterized in this experimental context. Excretion is regulated by interacting autonomic, enteric, and central mechanisms, and NMDA receptor blockade may influence these systems at multiple levels. MK-801 has previously been reported to alter respiratory function.^84^ Nevertheless, because MK-801 was administered systemically and urine output was estimated semi-quantitatively from the change in filter-paper mass, the present results do not identify the physiological mechanism underlying the reduction in excretion. Direct physiological measurements and central versus peripherally restricted manipulations will be required to clarify whether these effects arise from central autonomic regulation, peripheral NMDA receptor signaling, changes in movement or fluid balance, or a combination of these factors.

Our study has several important limitations. First, the recording system did not permit individual USVs to be assigned to either member of the male–female dyad. Because males are generally the primary vocalizers during male–female interactions, the observed reduction most likely reflected suppression of vocal production by the MK-801-treated male. However, a contribution from the untreated female cannot be completely excluded because our recording system did not permit caller identification. Methods that enable caller identification will be required to quantify the relative vocal contributions of the male and female. Second, we examined only C57BL/6J mice, which limits the generalizability of our findings across mouse strains and species. Species-specific differences in the contexts and potential functions of USVs have been reported. For example, rats emit distinct USVs in both appetitive and aversive contexts, whereas comparably clear valence-related patterns have not been established in mice.^85^ Comparisons across mouse strains and with rats and other species would therefore be valuable for evaluating the generalizability and translational relevance of the present findings. Third, the estrous stage of the stimulus females was not determined. Previous findings regarding the influence of female reproductive status on courtship USVs have varied according to the experimental context and the vocal measure examined. Female estrous stage or sexual cycle has been reported to influence the acoustic characteristics and emission patterns of courtship USVs without necessarily altering the total number of calls, whereas ovariectomy did not significantly affect the total number or frequency-modulated subtypes of USVs recorded during male-female interactions in another study.^34,86,87^ Because estrous stage was not monitored in the present study, its potential contribution to variability in the dyadic vocal and interaction measures could not be evaluated. Fourth, systemic MK-801 administration did not allow us to distinguish among the neural, peripheral, behavioral, and physiological processes that may have contributed to the observed effects. NMDA receptors are expressed not only in the central nervous system but also in peripheral systems, including the enteric nervous and cardiovascular systems. Accordingly, the changes in locomotor activity, excretory outcomes, and dyadic USVs may have involved central NMDA receptor blockade, peripheral signaling, altered respiratory or motor control, changes in social interaction, or a combination of these factors. The present experiments were not designed to distinguish among these mechanisms. Finally, we administered MK-801 only to male mice and therefore did not examine its effects when the female member of the dyad was treated. The present findings cannot establish whether NMDA receptor blockade has sex-dependent effects on social behavior, dyadic vocal communication, locomotor activity, or autonomic and excretory function. Future studies incorporating caller-identification methods, estrous-stage monitoring, direct comparisons of MK-801 administration to males and females, cross-strain and cross-species designs, and region-specific or peripherally restricted pharmacological manipulations will be required to address these questions and clarify the mechanisms underlying the observed effects. In conclusion, systemic MK-801 administration to adult male mice dose-dependently reduced the number of USVs recorded during male–female interactions and produced distinct effects on social approach, locomotor activity, and excretory outcomes. These findings demonstrate that systemic NMDA receptor blockade differentially affects multiple behavioral domains but do not establish the neural mechanisms underlying the reduction in vocalizations.

## Glossary

NMDA: N-methyl-D-aspartate
USVs: ultrasonic vocalizations

## Data availability

The data supporting the findings of this study are available from the corresponding author upon reasonable request.

## Code availability

The original code for the analyses is available from the corresponding author upon reasonable request.

## Ethical statement

All animal procedures were approved by the Animal Research Committee of Keio University and were conducted in accordance with the institutional guidelines for the care and use of laboratory animals. The study is reported in accordance with the ARRIVE guidelines.

## Competing interests

The authors declare no competing interests.

## Funding

This work was supported by JSPS KAKENHI 18KK0070 (KT), 19H05316 (KT), 19K03385 (KT), 19H01769 (KT), 20J21568 (KY), 22H01105 (KT), 23H02787 (KT), 23K27478 (KT), 23K22376 (KT), 24H00729 (KT), 26K00589 (KT), 26K22216 (KT), 24K06626 (KH), 25KJ0306 (KH), 24K16869 (KY), and 24KJ0069 (KY), Keio Academic Development Fund (KT), Keio Gijuku Fukuzawa Memorial Fund for the Advancement of Education and Research (KT), the Smoking Research Foundation (KT), and HOKUTO Foundation for the Promotion of Biological Science (KT).

## Author Contributions (CRediT)

Conceptualization: RT, KT. Methodology: RT, KT, KY, DN, KH, MT, YU, HK, MY, SM. Investigation: RT, KT, KY, DN, KH, MY, SM. Resources: MT, YU, HK. Formal analysis: RT, YU, MY, KT. Visualization: RT, YU, MY, KT. Supervision: KT. Project administration: KT. Writing—original draft: KT. Writing—review and editing: KT. All authors reviewed and approved the final version of the manuscript.

## Acknowledgements

We thank Shohei Kaneko, Hiroto Inoue, Shunsuke Nakajima, and Yuta Tamai for their assistance with animal care and valuable discussions. We also sincerely thank the anonymous reviewers for their careful evaluation and constructive comments, which substantially improved the clarity, accuracy, and overall quality of the manuscript. During revision, the authors used ChatGPT (OpenAI) solely to assist with English-language editing and improve the clarity of the manuscript. All scientific content, interpretations, references, and statistical results were independently reviewed and verified by the authors, who take full responsibility for the final manuscript.

## Notes

### Competing Interest Statement

The authors have declared no competing interest.

### Summary of Updates

The manuscript was revised in response to editorial feedback. We clarified the framing of the Introduction by condensing the schizophrenia-related background and focusing the rationale directly on the effects of pharmacological NMDA receptor blockade on ultrasonic vocalizations and social approach in mice. The study data and principal conclusions remain unchanged.

